# JNK signaling regulates oviposition in the malaria vector *Anopheles gambiae*

**DOI:** 10.1101/2020.03.14.990945

**Authors:** Matthew J. Peirce, Sara N. Mitchell, Evdoxia G. Kakani, Paolo Scarpelli, Adam South, W. Robert Shaw, Kristine L. Werling, Paolo Gabrieli, Perrine Marcenac, Martina Bordoni, Vincenzo Talesa, Flaminia Catteruccia

## Abstract

The reproductive fitness of the *Anopheles gambiae* mosquito represents a promising target to prevent malaria transmission. The ecdysteroid hormone 20-hydroxyecdysone (20E), transferred from male to female during copulation, is key to *An. gambiae* reproductive success as it licenses females to oviposit eggs developed after blood feeding. Here we show that 20E-triggered oviposition in these mosquitoes is regulated by the stress- and immune-responsive c-Jun N-terminal kinase (JNK). The heads of mated females exhibit a transcriptional signature reminiscent of a JNK-dependent wounding response while mating — or injection of virgins with exogenous 20E — selectively activates JNK in the same tissue. RNAi-mediated depletion of JNK pathway components inhibits oviposition in mated females, whereas JNK activation by silencing the JNK phosphatase *puckered* induces egg laying in virgins. Together, these data identify JNK as a potential conduit linking stress responses and reproductive success in the most important vector of malaria.

## INTRODUCTION

*Anopheles gambiae* mosquitoes are the most important vectors for *Plasmodium* malaria parasites, which infected at least 200 million people and caused more than 400,000 deaths in 2018 (WHO, 2019). The number of malaria deaths has more than halved since the year 2000 largely as a result of mosquito control strategies, especially insecticide-treated bed nets (WHO, 2019). This promising progress is, however, threatened by the spread of insecticide resistance in *Anopheles* populations (Hemingway, 2014), highlighting the pressing need for novel strategies for mosquito control. A number of recently proposed alternatives aim at reducing vector populations by regulating female reproductive output via either chemical (Childs et al., 2016) or genetic (Hammond et al., 2016) means, and their successful development is dependent upon a detailed understanding of the mechanisms regulating reproduction in *Anopheles* (Mitchell and Catteruccia, 2017).

Mating represents a vulnerable step in the *An. gambiae* reproductive cycle as it happens only once in the female’s lifetime. During this single sexual event, males transfer sperm along with a gelatinous mating plug that contains a host of proteins and other factors produced by the male accessory glands (Baldini et al., 2012; Rogers et al., 2009), including the ecdysteroid hormone 20-hydroxyecdysone (20E) (Baldini et al., 2013; Pondeville et al., 2008). Sexual transfer of this steroid hormone is a feature that is unique to anophelines, having evolved specifically in the *Cellia* subgenus and exclusively in the lineages leading to the most important African and South East Asian malaria vectors (Mitchell et al., 2015; Pondeville et al., 2019). Transfer of 20E drives profound behavioral and physiological changes in the female collectively termed post-mating responses (Baldini et al., 2013; Gabrieli et al., 2014; Shaw et al., 2014). Perhaps the most striking of these changes are refractoriness to further mating, underpinning the female’s monandry, and a license to oviposit eggs developed following a blood meal (Gabrieli et al., 2014). Depletion of endogenous 20E levels in males reduces egg laying rates and increases remating frequency in the females with whom they mate (Gabrieli et al., 2014). These effects are recapitulated by injection of exogenous 20E in virgin females, which is sufficient to induce both oviposition of developed eggs and refractoriness to further copulation (Gabrieli et al., 2014). Importantly, both refractoriness to further mating and the license to oviposit are irreversible, lifelong behavioral switches. However, the molecular processes through which 20E induces these changes in *An. gambiae* remain unknown.

Some insight into the mechanisms regulating these processes may come from the distantly related dipteran model organism, *Drosophila melanogaster*, where — similar to *An. gambiae* — the effects of mating include both a temporary refractoriness to further copulation and increased oviposition. Interestingly, while 20E in *D. melanogaster* does have an important role in regulating courtship behavior and specifically in the consolidation of long-term courtship memory in males (Ishimoto and Kitamoto, 2011; Ishimoto et al., 2009), post-mating responses in fruit fly females are not driven by 20E but by small male accessory gland peptides (Acps) transferred to the female during mating. These include ovulin, which is implicated in the control of oviposition (Rubinstein and Wolfner, 2013), and most importantly Acp70A, also known as Sex Peptide (SP) (Chen et al., 1988). SP is necessary and sufficient to induce the post-mating switch: injection of exogenous SP induces both refractoriness to mating and oviposition (Chen et al., 1988) while loss of the G protein-coupled Sex Peptide Receptor (SPR) in females largely blocks these responses (Yapici et al., 2008). Consistent with its role in modulating female post-mating behavior, SPR is found in neurons innervating the reproductive tract as well as the brain and ventral nerve chord (Wang et al., 2020; Yapici et al., 2008), while ovulin acts through octopamine-dependent neurons (Rubinstein and Wolfner, 2013). The importance of the brain in controlling female responses after copulation in *Drosophila* is also demonstrated by the fact that in genetic SPR mutants post-mating responses can be rescued by introducing a mutation that yields a leaky blood-brain barrier phenotype (Haussmann et al., 2013), identifying the entry of mating factors into the brain as a potentially crucial step in inducing post-mating changes. Consistent with these findings, the *Drosophila* brain exhibits a robust transcriptional program following mating or injection of exogenous SP (Dalton et al., 2010).

Here we show that 20E-induced oviposition behavior in *An. gambiae* is partially regulated by c-Jun N-terminal kinase (JNK) signaling in the female head. We detect a strong, mating-induced transcriptional signature in female heads, enriched in immune genes and reminiscent of a JNK-dependent wound-healing response. Silencing multiple components of JNK signaling reduces oviposition rates of mated females, as well as those of virgin females injected with 20E. Conversely, JNK activation by depletion of the negative regulator *puckered* increases oviposition rates in virgin females. Our results unveil an unexpected link between an important mosquito reproductive behavior and the activation of JNK, a pathway classically associated with stress resistance and longevity (Wang et al., 2003, 2005) but which is also pivotal to anti-plasmodium immunity (Garver et al., 2013; Ramphul et al., 2015).

## RESULTS

### A transcriptional signature of wounding response is found in the head after mating

To gain insight into the molecular basis of the mating response in *An. gambiae*, we performed transcriptional analysis of the heads of mated and age-matched virgin females at 3 and 24 hours post mating (hpm). Microarray analysis identified a specific immune signature triggered by mating in the head, the like of which had not been detected in similar analyses of other *An. gambiae* female tissues (Gabrieli et al., 2014; Rogers et al., 2008; Shaw et al., 2014). As summarized in Table 1, 23 genes were differentially regulated after mating at the two time points under analysis, 22 of which were upregulated at either 3 hpm (11 genes) or 24 hpm (11 genes). A single gene, an acyltransferase (AGAP007078), was down regulated in the head 24 hpm. Functionally, 6 of the 22 upregulated genes were common to a group of genes previously linked to the wounding response in *An. gambiae* (Nsango et al., 2013), while 16 were implicated in the melanization pathway, which has been studied in this species predominantly in the context of *Plasmodium* infection (Barillas-Mury, 2007; Michel and Kafatos, 2005) but is also strongly linked to wound healing in other insects including *Drosophila* (Bidla et al., 2009; Lee and Miura, 2014). In addition, 13 of the upregulated genes were previously found to be preferentially expressed in hemocytes (Pinto et al., 2009), cells related to mammalian macrophages and central to the repair and regeneration of damaged cells (Fogarty et al., 2016) and to the wounding-induced transcriptional response in *Drosophila* (Stramer et al., 2008).

**Table 1.**
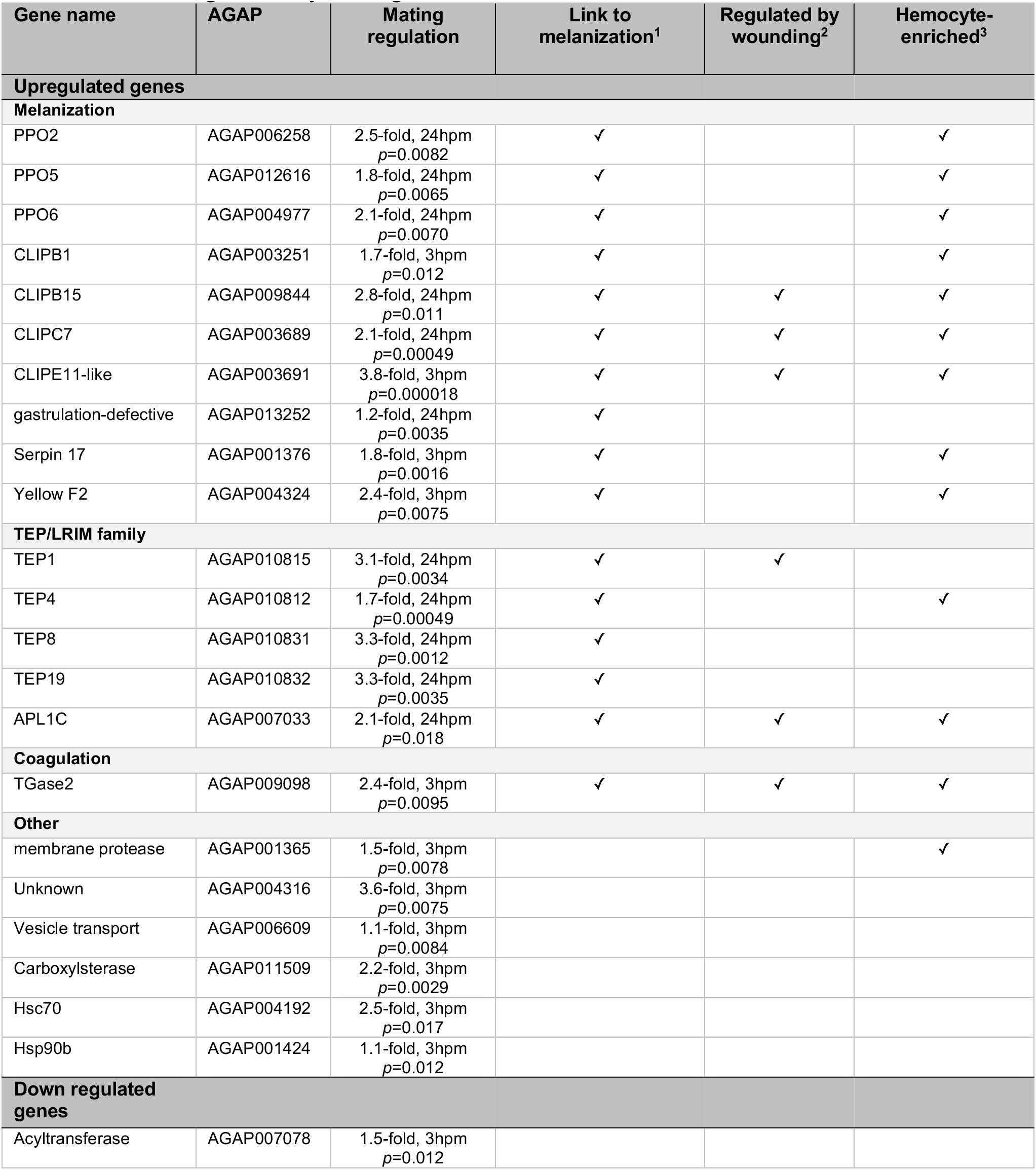
Genes regulated in the head after mating are enriched in genes linked to wound-healing, hemocytes and the JNK pathway. The 23 genes identified by microarray as being significantly upregulated in the head after mating were compared with relevant literature reports: ^1^ gene families linked to the melanization pathway (Barillas-Mury, 2007; Michel and Kafatos, 2005); ^2^ genes upregulated by wounding in *An. gambiae* (Nsango et al., 2013); ^3^ genes preferentially expressed in *An. gambiae* hemocytes (Pinto et al., 2009). The mean mating-induced fold-change over age-matched virgins, the time point at which that change was observed and the adjusted *p* value, after False Discovery Rate (FDR) correction for multiple testing, are indicated (further detail of statistical validation of microarray data is included in Methods).

At the earlier 3 hpm time point we found mating-induced upregulation of transglutaminase 2 (TGase2), a member of an enzyme family involved in chemical crosslinking of proteins in the hemocel that is implicated in the coagulation responses that follow infection or trauma (Bidla et al., 2009) and was previously shown to be involved in JNK-dependent wounding responses in *An. gambiae* (Nsango et al., 2013) (Table 1). Also upregulated at 3 hpm were two members of the CLIP protease family, CLIPs B1 and 11E-like, serine proteases which initiate proteolytic cascades leading to the activation of prophenol oxidases (PPOs) that mediate both melanization (Christensen et al., 2005) and coagulation (Bidla et al., 2009). We also identified the CLIP inhibitor, Serpin 17, a member of the serine protease inhibitor (Serpin) family which block CLIP-mediated proteolytic processing of PPO enzymes (Tang, 2009). Finally, we detected the up-regulation of L-dopachrome tautomerase (also known as Yellow F2), an enzyme involved in melanin biosynthesis (Christensen et al., 2005) (Table 1).

Among the genes upregulated at 24 hpm we identified three PPOs (PPO2, PPO5, and PPO6) and three additional CLIP proteases; CLIP15B, CLIPC7, and *gastrulation defective*. Beyond the melanization pathway we also found multiple thioester-containing proteins (TEP1, TEP4, TEP8 and TEP19) which are linked to the mosquito complement-like response (Blandin et al., 2008), along with APL1C (*Anopheles-Plasmodium*-responsive leucine-rich repeat protein 1, isoform C), a member of the leucine-rich repeat (LRR) immune protein (LRIM) family. APL1C associates with another LRIM family member (LRIM1, not identified here), and TEP1 in a complex that retains TEP1 in a stably active form that can then form thioester bonds with surface-exposed proteins of invading pathogens including *Plasmodium* parasites (Blandin et al., 2004; Fraiture et al., 2009; Povelones et al., 2011), targeting them for lysis. Taken together, these data suggest mating induces a wounding response specific to the head of *An. gambiae* females.

### JNK is activated in the head after mating

The preponderance of wounding-related transcripts in the head after mating prompted us to ask what signaling pathways might lead to such a response. The importance of the JNK signaling pathway in the insect wounding response has been highlighted in both *An. gambiae* (Nsango et al., 2013) and *Drosophila* (Ramet et al., 2002). Since JNK is well known to be activated post-transcriptionally by phosphorylation, we asked whether mating might increase levels of active (phosphorylated) JNK (pJNK) in the female’s head. *An. gambiae* females were dissected around the onset of detectable transcriptional changes (1-6 hpm), and heads were analyzed by Western blot using an antibody specifically recognizing pJNK. Mating induced a marked increase in the levels of pJNK in the head compared to virgin controls, an effect that was already detectable at 1 hpm and was still evident at 4 hpm across multiple experiments (Fig. 1a, b). In mated females we detected a strong band at 46 kDa and a much weaker band at 52 kDa, similar to reports in *An. stephensi* (Souvannaseng et al., 2018). This mating-induced JNK response was tissue-specific as we did not detect it in other tissues including the reproductive tract (ovaries, atrium and spermatheca) (Fig. 1c, Fig. S1a, b). Moreover, while levels of pJNK increased in the heads of mated females (one sample t test, p=0.0496, n=4), levels of the phosphorylated (active) form of the related MAP kinase extracellular signal-regulated kinase (pERK) were reduced in mated heads relative to virgin controls (one sample t test, *p*=0.02, n=4) (Fig. S1c), suggesting some level of pathway specificity.

**Figure 1.**
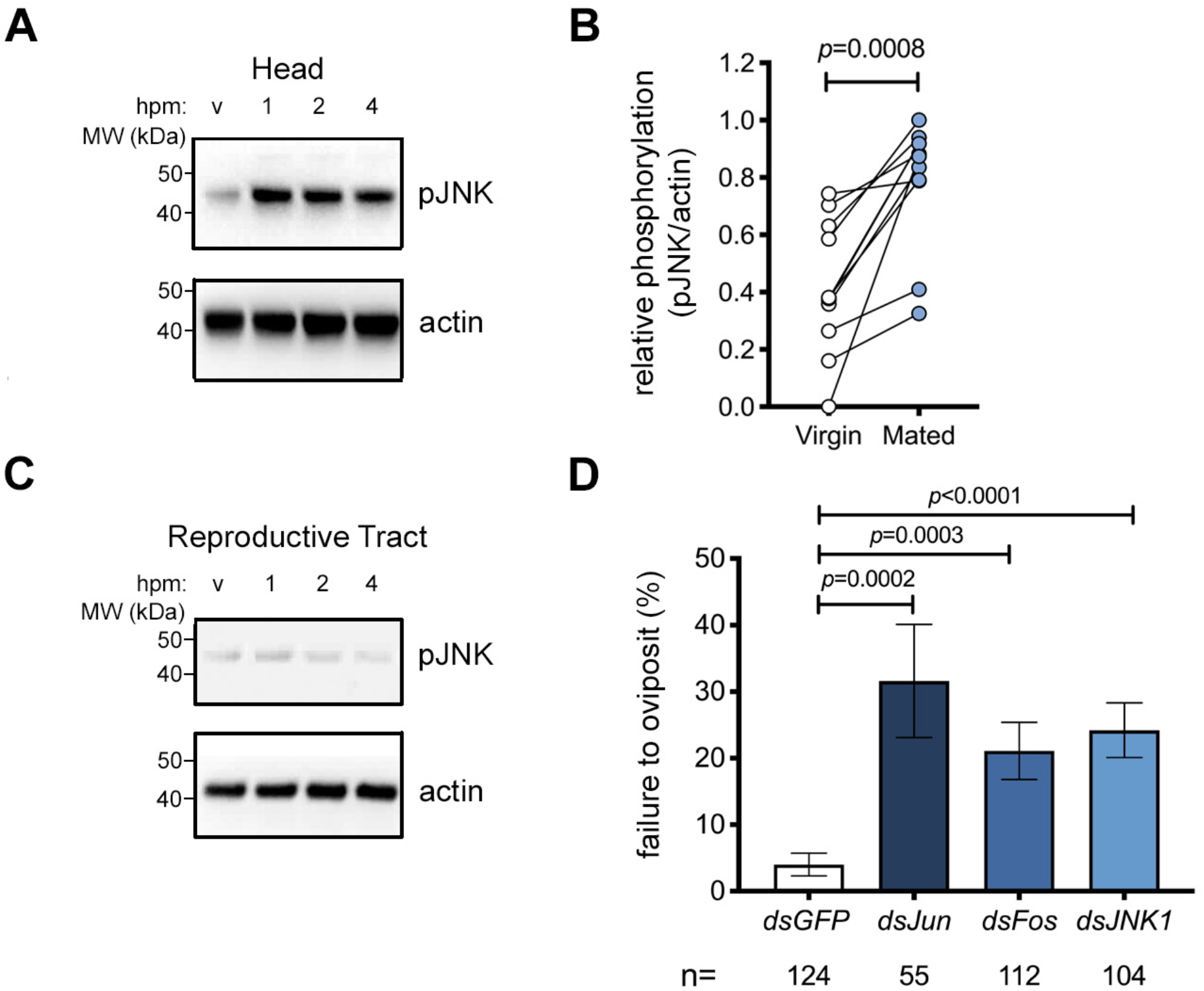
JNK is activated in the head after mating and required for mating-induced oviposition. **A-C.** Representative Western blot of extracts of **(A)** heads or **(C)** reproductive tracts (ovaries, atrium and spermatheca) prepared from virgin or mated females at 1, 2 and 4 hours post mating (hpm). Tissue extracts were subjected to Western blot analysis with anti-pJNK then stripped and re-probed with anti-actin as loading control. In **B,** the optical density of bands was quantified (ImageJ) in different experiments, and the pJNK signal was normalized against actin and expressed as ‘relative phosphorylation’ in virgins and females 2 hpm. Differences in the relative pJNK levels in virgin and mated females were analyzed using a Mann-Whitney test and significant *p* values (*p*<0.05) reported. **D.** RNAi silencing of JNK (ds*JNK*), Jun (ds*Jun*) or Fos (ds*Fos*) prior to mating reduces the oviposition rates of mated females. The graph shows the percentage of females (mean ± SEM from 9 independent biological replicates) failing to oviposit by day 4 post mating, analyzed using a logistic regression test. For the dataset as a whole, chi-squared=49.1, *p*<0.0001.

### JNK1 is required for mating-induced oviposition

To determine the contribution of the JNK pathway to post-mating responses, we silenced elements of the pathway by RNAi and examined the effect on oviposition, a physiological response that is induced in mated females once eggs are fully developed after blood feeding. Given the annotation of two distinct *An. gambiae JNK* genes, *JNK1* (AGAP029555) and *JNK3* (AGAP009460), we performed preliminary experiments to assess the relative expression of each transcript and found that *JNK1* transcript levels were 2–3 orders of magnitude more abundant than *JNK3* in all tissues measured (Fig. S2) leading us to focus on the *JNK1* gene product. Thus, virgin females were injected with ds*RNA*s targeting *JNK1* (*dsJNK1*, Fig. S3) or its two transcription factor targets, *Jun* (ds*Jun*) and *Fos* (ds*Fos*), using ds*GFP* as control. Using a previously established protocol (Gabrieli et al., 2014), injected females were then blood-fed and mated, and oviposition rates were measured. *JNK1* silencing inhibited the mating-induced increase of both strong (46kDa) and weak (52kDa) pJNK bands in the head (Fig. S4) and significantly prevented oviposition compared to control females (logistic regression model, *p*<0.0001, Fig. 1d). The failure to oviposit after mating was 5.6-fold more likely in *dsJNK1*-treated females than in *dsGFP*-treated controls (odds ratio [OR]=5.6; *p*<0.0001). Similarly increased rates of oviposition failure were observed with *dsFos* (OR=4.03, *p*=0.0003) and ds*Jun* (OR=6.2, *p*=0.0002) (Fig. 1d) while we observed no significant effect on the total number of eggs developed (Fig. S5a) or the number of eggs oviposited (Fig. S5b) in any group. Moreover, *JNK1* depletion reduced the up-regulation of wounding-related genes (*APL1C, TEP1* and *PPO2*) in the female head after mating (Fig. S6). Together, these data suggest the involvement of the JNK pathway in the mating-induced cascades leading to oviposition in *An. gambiae*.

### Depletion of the JNK-phosphatase puckered is sufficient to induce oviposition in virgins

Having established a role for the JNK pathway in mating-induced oviposition, we went on to determine whether an increase in the level of pJNK in the head might be sufficient, *per se*, to induce oviposition in blood-fed virgin females. The activation of JNK is regulated by the dual phosphorylation of its TxY motif by the MAP kinase kinase (MAP2K) Hemipterous, and is prevented by dephosphorylation of the same motif by the dual specificity phosphatase Puckered (Puc), also known as MAP kinase phosphatase 5 (MKP5). Because depletion of *puc* in *Drosophila* leads to the spontaneous activation of JNK-responsive genes (Martin-Blanco et al., 1998), we reasoned that silencing of this phosphatase might mimic the activation of JNK noted after mating. Indeed, *puc* silencing (Fig. S3) increased pJNK levels relative to controls in the female head across multiple experiments (Fig. 2a, b), while no noticeable effects were observed in the reproductive tract of the same females suggesting tissue-specific activation (Fig. 2c). Moreover, a significant proportion of virgin females (logistic regression model, *p*<0.0001) injected with *dspuc* laid eggs after blood feeding (Fig. 2d). Using the logistic regression model applied above, we determined that ds*puc* virgins were approximately 25-fold more likely to oviposit than controls (OR=24.6, *p*<0.0028). These results are consistent with our findings that JNK activation in the head after mating induces oviposition.

**Figure 2.**
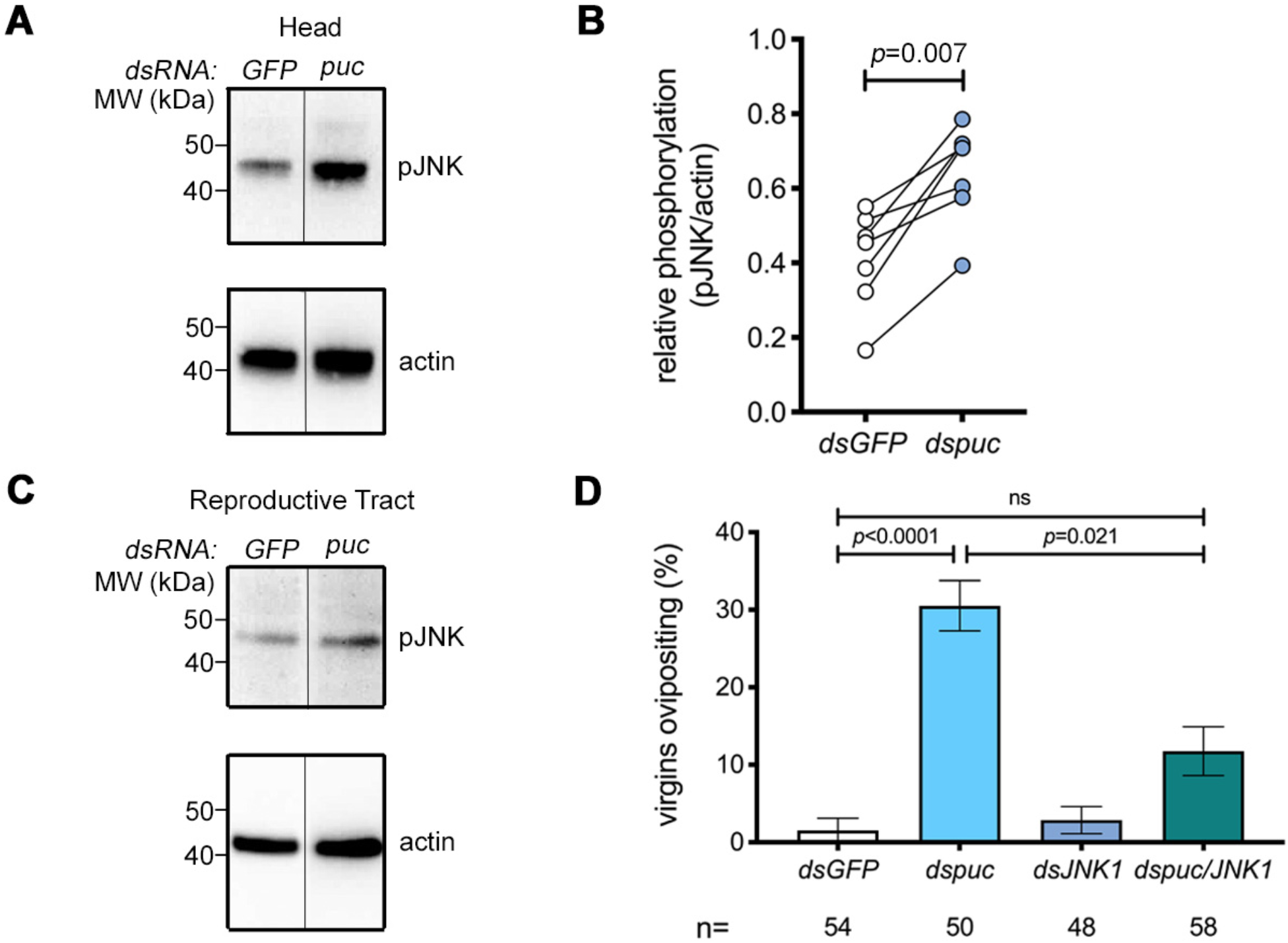
*Puckered* knock down induces phospho-JNK in the head and JNK1-dependent oviposition in blood-fed virgins. **A-C.** Representative Western blot of extracts of **(A)** heads or **(C)** reproductive tracts (ovaries, atrium and spermatheca) dissected from virgin or mated females injected with either *dsGFP* (*GFP*) or *dspuc* (*puc*) 48 hours post injection (hpi), using anti-pJNK then stripped and re-probed with anti-actin as loading control. In **B,** the optical density of bands was quantified (ImageJ) in different experiments, and the pJNK signal was normalized against actin and expressed as ‘relative phosphorylation’ of pJNK in *dsGFP*- and *dspuc*-treated females. The line between *GFP* and *puc* indicates the removal of an unrelated intervening lane. Differences in the relative pJNK levels in *dsGFP* and *dspuc*-treated females were analyzed using a Mann-Whitney test and significant *p* values (*p*<0.05) reported. **D.** RNAi silencing of *puc* (*dspuc*) induces oviposition in blood fed virgins. Virgin females were injected with *dsGFP, dsJNK, dspuc* or jointly injected with *dsJNK and dspuc*, blood-fed and then placed in oviposition cups. The graph shows the percentage of females (mean ± SEM from 4 independent biological replicates) successfully ovipositing by day 5 post blood feeding analyzed using a logistic regression test. For the dataset as a whole chi-squared=28.4, *p*<0.0001.

While in *Drosophila* Puc acts selectively to regulate the JNK pathway (Martin-Blanco et al., 1998), a recent study in *An. stephensi* suggested that the same phosphatase can impact MAP kinases other than JNK (Souvannaseng et al., 2018). To address the role of JNK1 in the oviposition induced by *dspuc*, we performed a double knock down of *puc* and *JNK1*. Consistent with a predominant role for JNK signaling in regulating this behavior, coinjection of *dspuc* with *dsJNK1* significantly reduced the frequency of oviposition induced by *dspuc* alone (OR relative to *dspuc* alone 3.7, *p*=0.021) without affecting the efficiency of *puc* knock down (Fig. S3). These data suggest that activation of JNK in the head of a virgin female is sufficient to induce oviposition.

### The JNK pathway is required for oviposition induced by exogenous 20E

Since thoracic delivery of exogenous 20E is sufficient to induce oviposition in blood-fed virgin females (Gabrieli et al., 2014), we next assessed whether injected 20E—like mating (Fig. 1 a,b)—might also induce an increase in pJNK levels in the head. When compared to controls, a robust increase in pJNK in the head was observed across multiple experiments 1–2 hours post injection (hpi) (Fig. 3a, b), while the reproductive tract again showed no JNK activation at these time points (Fig. 3c).

**Figure 3.**
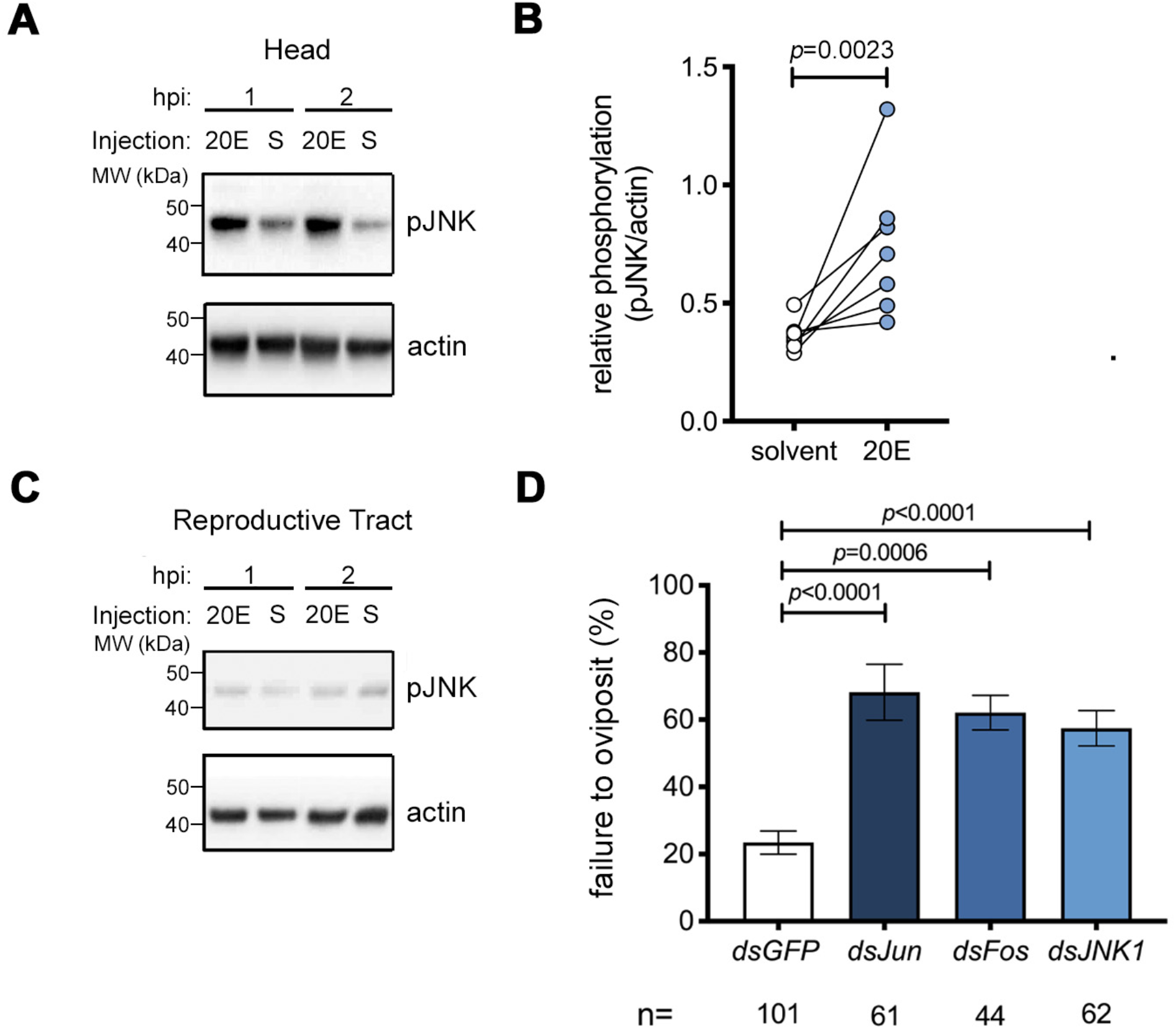
JNK pathway depletion causes failure of 20E-induced oviposition in blood-fed virgins. **A-C.** Representative Western blot of extracts of **(A)** heads or **(C)** reproductive tracts (ovaries, atrium and spermatheca) dissected from virgin females injected with either 20E or a solvent control (S) at 1 or 2 hours post injection (hpi) using anti-pJNK, and anti-actin as loading control. In **B,** the optical density of bands was quantified (ImageJ) in different experiments, and the pJNK signal was normalized against actin and expressed as ‘relative phosphorylation’ of pJNK in females injected with solvent or 20E (2hpi), analyzed using a Mann-Whitney test. **D.** Virgin females were injected with ds*JNK*, ds*Jun* or ds*Fos*, blood-fed and then injected with 20E or solvent control and placed in oviposition cups. The graph shows the percentage of females (mean ± SEM from 7 independent biological replicates) failing to oviposit by 4 days post 20E injection. Differences in the relative likelihood of failed oviposition between the indicated groups were analyzed using a logistic regression test. For the dataset as a whole chi-squared=43.5, *p*<0.0001.

To determine if 20E-induced oviposition occurs via JNK signaling, we silenced *JNK1*, *Jun* or *Fos* and allowed females to take a blood meal. After completion of egg development, females were injected with either 20E or a solvent control, and oviposition rates were measured. The number of females failing to oviposit after 20E-injection increased markedly after treatment with *dsJNK1* (logistic regression model, *p*<0.0001), compared to *dsGFP*-treated control females (OR of failed oviposition compared to *dsGFP* control=5.4, *p*<0.0001). Similar results were obtained after silencing of *Jun* (OR 5.8, *p*<0.0001) and *Fos* (OR 4.1, *p*=0.0006) (Fig. 3d). These data support the involvement of the JNK pathway in oviposition induced by sexual transfer of the male steroid hormone 20E. Injection of a solvent control failed to induce oviposition in any ds*RNA* group (94-100% failed oviposition, see Table S1), as expected (Gabrieli et al., 2014).

## DISCUSSION

The transfer of the ecdysteroid hormone 20E from *An. gambiae* males to females during copulation is linked to a life-long license to oviposit eggs developed across multiple gonotrophic cycles (Gabrieli et al., 2014; Mitchell et al., 2015). Here, we identify the JNK pathway, traditionally associated with responses to diverse environmental stressors including wounding (Nsango et al., 2013) and *Plasmodium* infection (Garver et al., 2013; Ramphul et al., 2015), as a key component of the downstream events linking sexual transfer of 20E to oviposition. We document a rapid and selective increase in the amount of active pJNK in the heads of mated females that is recapitulated by injection of exogenous 20E. The activation of JNK in the head after mating appears specific given the absence of activation of the related MAP kinase, ERK, in the same tissue or of JNK itself in other tissues (reproductive tract or rest of body). This organ specificity is consistent with our previous microarray analyses that revealed no mating-induced wound-healing and immune responses in other female tissues, despite a strong transcriptional response involving hundreds of genes (Gabrieli et al., 2014; Rogers et al., 2008; Shaw et al., 2014), and contrasts with the striking immune signature and copulatory wounding reported in *Drosophila* reproductive tract after mating (Mack et al., 2006) (Mattei et al., 2015).

We find that the activation of the JNK pathway in the head is both necessary and sufficient for at least one element of the post-mating response - the oviposition of developed eggs. RNAi-mediated depletion of *JNK1* inhibited the mating-induced pJNK signal (Fig. S4) and reduced oviposition induced either by mating (Fig. 1d) or by the injection of exogenous 20E (Fig. 3d). The link between this reduced egg laying phenotype and JNK function is highlighted by the fact it was phenocopied by depletion of *Jun* or *Fos*, transcription factor targets of pJNK. Moreover, activation of the JNK pathway — by RNAi-induced depletion of *puc —* was sufficient to both increase pJNK levels in the head (Fig. 2a, b) and trigger oviposition in blood-fed-virgin females (Fig. 2d). Importantly, while recent data have shown that in some settings puc is capable of modulating MAP kinase pathways other than JNK (Souvannaseng et al., 2018), our data show that the induction of oviposition by depletion of *puc* is largely reversed by concomitant depletion of JNK1, identifying this MAP kinase as the dominant puc target in regulating this phenotype (Fig. 2d).

Despite the 100-1000-fold greater transcript abundance of *JNK1 (*Fig. S2), we cannot exclude a contribution to the oviposition phenotype from the *JNK3* gene product, given the 70% identity between the two over the mRNA sequence targeted by *dsJNK1*. Similarly, the origin and importance of the two protein bands observed using the pJNK antibody remain unclear. They could reflect the products of the *JNK1* and *JNK3* genes as suggested by others (Souvannaseng et al., 2018) but might equally represent alternative post-translationally modified forms of the same gene product. Future studies using specific antibodies or gene targeting strategies will help address these questions.

Given the partial nature of the effects observed, our data also highlight the likely existence of JNK-independent pathways controlling oviposition, although we cannot exclude that these effects are due to incomplete gene silencing. Importantly, in the absence of tissue-specific gene knockout studies, the systemic nature of RNAi makes it difficult to exclude the possibility that JNK signaling outside the head may contribute to the regulation of oviposition, notwithstanding the head-specific activation of JNK observed after mating, 20E injection or puc depletion discussed above. The effects of the JNK pathway appear to affect oviposition rather than oogenesis, since neither the number of eggs developed or laid are affected by JNK pathway depletion (Fig. S5a, b).

While, to our knowledge, this is the first demonstration that the stress-responsive JNK pathway is involved in an important reproductive behavior such as oviposition, egg laying has been previously linked to stress responses in mosquitoes. Stressful stimuli including heat, dessication, starvation and infection have all been shown to impact on the timing of oviposition (Canyon et al., 1999; Shaw et al., 2016; Sylvestre et al., 2013), whereas confinement stress has been hypothesized to increase the frequency of oviposition in various anophelines (Nepomichene et al., 2017). Links between steroid hormones and stress responses have also been documented (Hirashima et al., 2000; Ishimoto and Kitamoto, 2010; Zheng et al., 2018). Of particular relevance, 20E titers have been shown to increase following stressful social interactions such as courtship (Ishimoto et al., 2009). This has led to the proposal that — akin to related sex steroids in mammals (Parducz et al., 2006) —20E functions in insects to consolidate stress-associated memory and to drive pathways of neuronal remodeling underpinning the development of appropriate adaptive behaviors (Ishimoto and Kitamoto, 2011).

Although the mating-mediated mechanisms controlling egg laying remain largely elusive, the data presented here provide clear evidence of collaboration between 20E and the JNK pathway to induce oviposition. In fact, such collaboration is well documented during larval/pupal metamorphosis in *Drosophila*, where 20E and the JNK pathway together drive waves of apoptosis which control the reshaping or destruction of obsolete larval tissues (Lehmann et al., 2002) as well as the remodeling of neuronal connections in the larval brain by axon and dendrite pruning (Zhu et al., 2019). It is plausible that an irreversible process of this type might regulate the lifelong behavioral changes induced by mating in *An. gambiae.* In future studies, it will be important to examine whether other elements of the post mating response, such as mating refractoriness, show a similar requirement for the JNK pathway. In addition, given the importance of the JNK pathway in the *An. gambiae* immune response to some *Plasmodium* infections (Garver et al., 2013; Ramphul et al., 2015) and the centrality of 20E to mosquito physiology and parasite development (Werling et al., 2019), it will be informative to examine whether 20E signaling in other contexts (e.g. after blood feeding) also engages with the JNK pathway.

## ACKNOWLEDGEMENTS

The authors gratefully acknowledge the collaboration of the Servizio Immunotrasfusionale, Ospedale Santa Maria della Misericordia di Perugia for provision of blood samples. The authors thank Dr Roberta Spaccapelo (Universita degli studi di Perugia) and Dr. Tony Nolan (Liverpool School of Tropical Medicine) as well as members of the F.C. lab for helpful suggestions and critical reading of the manuscript. This study was sponsored by European Research Council 7th Research Framework Programme (Project ‘Anorep’ Starting Grant 260897), by a Faculty Research Scholar Award by the Howard Hughes Medical Institute and the Bill & Melinda Gates Foundation (Grant ID: OPP1158190), and by the National Institutes of Health (R01 AI124165, R01 AI104956) to F.C..

## Author Contributions

Conceptualization, M.P., S.N.M. and F.C; Methodology, M.P., S.N.M., E.K., P.G., W.R.S., F.C.; Investigation, M.P., S.M., E.K., P.S., P.G., K.W., P.M., P.G. and M.B.; Formal analysis, M.P., S.N.M., E.K., A.S., K.W., P.M., W.R.S., P.S. and P.G.; Writing-Original Draft, M.P. and F.C.; Review and editing, M.P., S.N.M., P.G., K.W., V.T. and F.C.; Visualization, M.P, S.N.M and F.C.; Funding Acquisition, F.C.; Resources, V.T. and F.C.; Supervision, M.P. and F.C.

## STAR METHODS

### KEY RESOURCE TABLE

STAR METHODS KEY RESOURCE TABLE
| REAGENT OR RESOURCE | SOURCE | IDENTIFIER |
| --- | --- | --- |
| **Antibodies** |  |  |
| Rabbit polyclonal anti-pJNK | Cell Signaling Inc. | #4668; RRID:AB_823588 |
| Rabbit polyclonal anti-pERK | Cell Signaling Inc | #9101; RRID:AB_331646 |
| Rat monoclonal anti-β actin (MAC237) | AbCam | Ab50591; RRID:AB_867488 |
| Goat anti-rabbit-HRP | Thermo Fisher | G-21234; RRID:AB_2536530 |
| Goat anti-rat-HRP | Thermo Fisher | 31470; RRID:AB_228356 |
| **Bacteria** |  |  |
| Top10 OneShot chemically competent *Escherichia Coli* | Thermo Fisher | C404003 |
| **Chemicals** |  |  |
| 20-hydroxyecdysone (20E) | Sigma Aldrich | H5142-5mg |
| **Critical Commercial Assays** |  |  |
| T7 MegaScript in vitro transcription kit | Thermo Fisher | AM1334 |
| Low input Quick Amp labelling kit | Agilent | 5190-2305 |
| 44-K *An. gambiae* whole genome microarray | Agilent | G2519F-020449 |
| **Deposited data** |  |  |
| Mating-regulated genes in head microarray | This paper | ArrayExpress<br># E-MTAB-8733 |
| Raw data used to prepare figures | This paper | Mendeley Data doi:<br>https://data.mendeley.com/data-sets/5gmsnv8zw9/draft?a=bc216e76-d54d-4744-98a6-5cd0cf60c8c5 |
| **Experimental Models** |  |  |
| Mosquito: *Anopheles gambiae* G3 | B.E.I Resources | MRA-112 |
| **Oligonucleotides** |  |  |
| Primers for qRT-PCR or dsRNA template production (See Table S2) | This paper | N/A |
| **Recombinant DNA** |  |  |
| pCR2.1Topo | Thermo Fisher | 450641 |
| pCR2.1 EGFP | Baldini et al, 2013 | N/A |
| pCR2.1 JNK1 | This paper | Adgene #133284 |
| pCR2.1 puc | This paper | Adgene #133285 |
| pCR2.1 Jun | This paper | Adgene #133286 |
| pCR2.1 Fos | This paper | Adgene #133287 |
| **Software and Algorithms** |  |  |
| Image J image analysis | Schneider et al, 2012 | https://imagej.nih.gov |
| E-RNAi design |  | https://www.dkfz.de/signaling/e-rnai3/ |
| JMP Pro13 | SAS | https://www.jmp.com/en_be/software/predictive-analytics-software.html |
| Prism 8 statistical and graphing software | Graph Pad | https://www.graphpad.com |

## EXPERIMENTAL MODEL AND SUBJECT DETAILS

### Rearing of *Anopheles gambiae* Mosquitoes

*An. gambiae* mosquitoes of the G3 line were maintained at 28°C, 70% relative humidity with a 12 hour light/ dark diurnal cycle and water and 10% glucose solution *ad libitum* and fed weekly on human blood (Servizio Immunotrasfusionale, Ospedale Santa Maria della Misericordia di Perugia). For mating experiments, males and females were maintained as virgins by separation of male and female pupae by microscopic examination of the terminalia and kept in separate cages until sexual maturity (3 days post eclosion).

## METHOD DETAILS

### Oviposition experiments

Sexually mature, three-day old virgin females were injected (Nanoject II, Drummond Scientific/Olinto Martelli Srl, Florence, Italy) with 0.69μg (138nl, 5μg/μl double stranded RNAs [dsRNAs, see below]) targeting JNK pathway components (*JNK1, Jun, Fos*, or *puckered [puc]*) or *GFP*, a gene not expressed in these mosquitoes, as a negative control. In experiments in which two genes were targeted simultaneously (e.g. Figure 2), a mixture of the two *dsRNAs,* in which the concentration of each was 5μg/μl, was prepared and injected in the same volume (138nl) as a single dsRNA treatment. Assignment of mosquitoes to different groups was random but groups were not blinded. The effect of gene depletion on oviposition was assessed essentially as previously described (Gabrieli et al., 2014). Three days post-injection, females were blood-fed and unfed females removed from the cage. Two days later (upon completion of egg development), blood-fed females were induced to oviposit developed eggs either by allowing them to mate (see below) or by injection of exogenous 20E (Sigma, Milan, Italy; see below). For mating-induced oviposition, females captured *in copula* were checked using a fluorescence microscope for the presence of a correctly-positioned, auto-fluorescent mating plug to ensure successful copulation. Mated females were allowed to recover overnight then placed in individual oviposition cups as described previously (Gabrieli et al., 2014). For ovpiposition induced by exogenous 20E, two days post blood feeding, females were injected (138nl) with 20E (38mM, equivalent to 2.5ng) dissolved in H2O containing 10%EtOH and 5% DMSO or with a solvent control (the same diluent minus 20E) and allowed to recover overnight before being placed in individual oviposition cups. As previously shown (Gabrieli et al., 2014) oviposition in our hands routinely takes place 2 days after mating or 20E injection. Thus, in accordance with previous similar experiments, mated or injected females were checked every day for four days after blood feeding and oviposition was deemed to have occurred if a single egg was detected in the oviposition cup. Females who died before ovipositing were excluded from the analysis, as were any females who failed to develop eggs on the assumption that they had not taken a blood meal. Statistical significance of differences between groups in the frequency of oviposition were assessed using a generalized linear model based on the binary outcome ‘oviposition or no oviposition’ by day 4 post mating or 20E injection or day 5 post blood feeding in the case of *dspuc* treatment. ‘Experiment’ was included as a variable in the model and experiments statistically distinguishable from the others were removed as outliers. In the case of mating-induced oviposition this led to the removal of 2 of 11 experiments while in the case of *dspuc*- or 20E-induced oviposition no outliers were identified. The model was used to calculate the odds ratios (OR) of the relative likelihood of oviposition, or failed oviposition, in each group as well as the statistical significance of inter-group differences.

#### Microarray experiments and analysis

Samples for analysis following mating were prepared and analyzed as previously described (Gabrieli et al., 2014). Briefly, heads from 15 mated or age-matched virgin control females were dissected in to ice-cold PBS at 3h and 24h post mating and immediately transferred to Tri Reagent (Thermofisher, Loughborough, UK). Four independent biological replicates were performed on different generations of the same mosquito line (G3). Total RNA was recovered using standard phenol/chloroform extraction, followed by DNase-treatment to remove genomic DNA and quantified using a NanoDrop spectrophotometer (Thermofisher). A one-color labelling strategy was used: RNA (100ng) from each of the four replicates was labelled using a Low Input Quick Amp Labeling kit (Agilent, Stockport, UK) following protocol G4140-90040. Labelled RNA was hybridised to 44-K *An. gambiae* whole genome microarrays (Design ID G2519F-020449). Labelling, hybridization and scanning were performed by the Institute of Genetics and Molecular and Cellular Biology (Illkirch, France).

Microarray datasets were analyzed using the R statistical software environment (version 2.15.0) running the Linear Models for Microarray Data (Limma) package (version 3.14.4) (Smyth et al., 2005). Single-color dye signals were background-corrected using the normexp method with an offset of 16 and were normalized across microarrays using the quantile method. Multiple probes for the same transcript identifier were collapsed to individual genes, and average fold-change results were generated for each unique array identifier. Package ArrayQualityMetrics (Kauffmann et al., 2009) was used for quality control of all microarrays before significance was estimated by fitting a linear model to each gene across replicated arrays, applying a contrast matrix of comparisons of interest in the mating-array experiment, and determining an empirical Bayes moderated t statistic. The decideTests function for multiple testing across genes and contrasts (“global strategy”) was also used to classify the related t statistics as up, down, or not significantly regulated. *P* values then were corrected for multiple testing by the Benjamini–Hochberg method (Benjamini and Hochberg, 1995). Results were exported as Microsoft Excel files, and transcript identifiers [Ensembl Gene ID (AGAP) numbers or ESTs] with adjusted P < 0.05 were selected for further analysis. When possible, ESTs were identified by using manual submissions of Blastn (Altschul et al., 1990) to the *An. gambiae* PEST strain genome or were classified as unknown. Gene function annotation was assigned via VectorBase gene description, AnoXcel summary (Ribeiro et al., 2004), and/or orthology.

### Western blotting

The heads, reproductive tracts (containing ovaries, atrium and spermatheca) and, on occasion, the rest of the body (RoB) were dissected from females (10-15 per point) treated as indicated in the text and placed in 25μl of homogenization buffer (Tris-HCl, 10mM (pH 7.4); NaCl, 150mM; EDTA, 5mM; Triton X-100 0.5%; sodium dodecylsulfate (SDS), 0.1% w/v; sodium orthovanadate 10mM, Sodium iodoacetate, 5mM; protease inhibitor cocktail (Sigma, prod. code, p8340) 10μl /ml; phosphatase inhibitor cocktail III (Sigma, prod. code, p0044) 10μl/ml; sodium fluoride (1mM) and frozen. Tissues were thawed and homogenized manually using a plastic pestle then debris pelleted (14000xg, 5 min, RT). The supernatant (18μl) was recovered and mixed with 6μl 4x LDS-PAGE sample buffer (Thermofisher) supplemented with dithiothreitol (DTT, 5mM) and denatured at 85°C for 5 min. Proteins were separated over 4-12% gradient Bis-Tris gels (Thermofisher) electrophoretically transferred to nitrocellulose membranes (Thermofisher) and blocked (1h, RT) in PBS-Tween (PBS containing 0.05% Tween-20 (Euroclone, Milan, Italy)) supplemented with 5% w/v bovine serum albumin (BSA) (anti-pJNK) or with 5% w/v non-fat milk (actin). Blocked membranes were incubated (overnight, 4°C, 1/1000 dilution in 5% BSA) with rabbit anti-pJNK (Cell Signaling Inc, Leiden, Netherlands, prod. code, 4668) then washed (4×15 min PBS-tween) and incubated (1h, RT, 1/5000 dilution in PBS-Tween containing 5% non-fat milk) with anti-rabbit HRP-conjugated secondary then washed again (4×15min, PBS-Tween) and developed using enhanced chemiluminescence (ECL) reagents (Ammersham, Cambridge, UK) and visualised using a Fusion FX chemiluminscence detector (Vilber-Lourmat, Marne-la-Vallée, France). Membranes were then stripped (15 min, RT, ReStore; Thermofisher) and re-probed (1h, RT, 1/10000 diluted in PBS-tween with 5% non-fat milk) with rat anti-b-actin (AbCam, Cambridge, UK, prod. code, Ab50591) washed (as above) and incubated with anti-rat-HRP (1h, RT, 1/5000 dilution in PBS-tween containing 5% non-fat milk) then washed and developed with ECL as above. Using exposures which led to protein signals defined by the Fusion FX software as ‘non-saturated’ the intensity of developed bands was analyzed using Image J software and above-background values for pJNK optical density were divided by the above-background value for actin in the same lane. This ratio (pJNK/actin) was used as a measure of ‘relative phosphorylation’ for each sample. Routinely, statistical significance of differences in relative phosphorylation between groups, was assessed by comparing the relative phosphorylation values in individual experiments in control vs test groups (virgin vs mated; *dsGFP* vs *dspuc;* solvent vs 20E), using an unpaired, two-tailed Mann-Whitney test. To compare the effect of mating on levels of pJNK and pERK in the same head samples, relative phosphorylation levels of each kinase in mated heads was divided by the value of the same kinase in virgin heads. These ‘fold-change’ values were compared to a hypothetical value of 1 using a one sample t-test which tested the null hypothesis that relative phosphorylation values in virgin and mated heads were equal. The t statistic and degrees for freedom were: pJNK, t=3.22, df=3; pERK, t=4.51, df=3. Both groups passed a Shapiro-Wilk normality test (pJNK, *p*=0.77; pERK, *p*=0.59).

### RNAi

Double-stranded RNA (dsRNA) constructs targeting *puckered* (*puc*, AGAP004353), *JNK1* (AGAP022950), *Jun* (AGAP006386) and *Fos* (AGAP001093) were prepared using established methods described elsewhere (Gabrieli et al., 2014). Briefly, PCR primers (see Table S2) specific to the gene of interest were designed using the E-RNAi webservice (https://www.dkfz.de/signaling/e-rnai3/) and used to generate blunt ended amplicons from *An. gambiae* cDNA which were ligated in to the TOPO 2.1 vector (Thermofisher) and transformed in to Top10 competent E Coli (Thermofisher) by heat shock. The purified plasmid was prepared by midiprep kit (Thermofisher) from blue/ white selected colonies and the insert verified by sequencing. These plasmids have been submitted to Adgene (pCR2.1 JNK1, #133284, pCR2.1 puc, #133285; pCR2.1 Jun, #133286, pCR2.1 Fos, #133287). These plasmids were used as a template to generate amplicons of each insert containing T7 RNA polymerase binding sites from which gene-specific dsRNAs were prepared using a T7 polymerase *in vitro* transcription kit (T7 Megascript, Thermofisher). DNase1-treated dsRNAs were purified by phenol/chloroform extraction, washed in ethanol and re-suspended in H2O at 10-20μg/μL. As a negative control a dsRNA targeting GFP was prepared in the same way from an EGFP-containing plasmid described previously (Marois et al., 2012). Experiments in which the efficiency of knockdown of the targeted gene was found to be less than 20% relative to levels in the *dsGFP* control were excluded from all analyses.

### Gene expression analysis by qRT-PCR

Knock down of targeted genes and mating-regulated expression of genes of interest was assessed using qRT-PCR using standard methods as described previously (Shaw et al., 2014). Briefly, dissected tissues were recovered to 10μl of RNA-later (Ambion) that was immediately supplemented with 250μl Tri-reagent (Thermofisher) and frozen. Samples were thawed, homogenized using a motorized pestle, and pestles washed with a further 100μl of Tri reagent. Homogenates were then centrifuged (14000 x g, 15 min, 4°C) and supernatants (300μl) mixed with an equal volume of 100% ethanol and transferred to Direct-zol RNA miniprep columns (Zymo Research/ Euroclone, Milan, Italy). RNA was washed, DNase-digested on the column and washed again then eluted in to 20μL H_2_O and RNA content measured spectrophotometrically. Some of this material (0.5-1μg) was reverse transcribed to cDNA in a reaction volume of 100μl as described in detail elsewhere (Gabrieli et al., 2014) and subsequently diluted with water to 250μl. Expression of genes of interest was measured in triplicate 5μl aliquots using Fast SybrGreen master Mix (Thermofisher) and the forward and reverse primers listed in Table S2 (all 300nM except *Rpl19* reverse primer which was used at 900nM). Reactions were run on a QuantStudio 3 thermocycler (Thermofisher). The number of cycles required to cross an automatically assigned threshold level of incorporated SybrGreen fluoresence (CT value) was calculated using the manufacturer’s software. For each sample, the mean CT value obtained for a reference gene (*Ribosomal protein L19*, *Rpl19*; AGAP004422) whose expression is insensitive to 20E-dependent stimuli used in these studies such as mating (Shaw et al., 2014) and blood feeding (Marinotti et al., 2005), was subtracted from the mean CT value for the gene of interest (delta CT). Samples lacking at least duplicate CT values separated by less than 0.5 cycles were rerun or discarded as unreliable. The amplification efficiency of all primer pairs used was found to be within the range 100±10%. Delta CT values were used to calculate the expression of the gene of interest relative to *Rpl19* using the formula: relative expression=2^(-delta CT). On occasion the fold-change in expression was calculated by dividing the relative expression of a gene of interest in extracts of control mosquitoes (e.g. virgins or *dsGFP*-treated females) by that in ‘treated’ mosquitoes (e.g. mated females, *dsJNK1*-treated) (delta delta CT). Effects of RNAi-mediated *JNK1* depletion on mating-induced expression changes were analyzed by 2-way ANOVA using relative expression data (2^-delta CT). Efficiency of knock down of target genes was calculated using delta delta CT values from *dstarget* vs *dsGFP* controls and analyzed statistically using a one sample t-test in which the null hypothesis tested was that the ratio of target expression between *dsGFP* to *dstarget* groups should be equal to 1. The t statistic and degrees for freedom for each group were: *dsFos* t=6.7, df=6; *dsJun* t=6.9, df=6; *dsJNK1* t=12.5, df=12; d*spuc* t=16.4, df=11; *dsJNK1* joint t=22.0, df=6; *dspuc* joint t=12.9, df=6. All groups passed the Kolmogorov-Smirnov normality test (*dsJNK1 [*p>0.1]; *dspuc* [*p*=0.058] *dsJun* [*p*>0.1]; *dsFos* [*p*>0.1]; *dsJNK1 joint,* [*p*>0.1]), *dspuc* joint (*p*>0.1).

## QUANTIFICATION AND STATISTICAL ANALYSIS

All Statistical analyses were performed using Prism 8.0 (GraphPad, La Jolla, USA) except logistic regression analyses which were performed using JMP version 14 (SAS, Cary, USA). Non-parametric methods were used unless the normality of all groups analyzed could be demonstrated using a Kolmogorov-Smirnov test. In all tests, a significance cut-off of *p*=0.05 was applied. Multiple comparisons of the same dataset were corrected using Dunnett’s (for parametric tests) or Dunn’s (for non-parametric tests) multiple comparison correction. On occasion (Figure S1c and Figure S2), the effect of a manipulation was assessed using a one sample t-test. The ratio of ‘test’ to ‘control’ values (test/control) was compared with a hypothetical value of 1, thus testing the null hypothesis that test and control values were equal. In these cases the normality of all groups was verified using a Kolmogorov-Smirnov test (Fig S2) or a Shapiro-Wilk test (Fig. S1c).

## DATA AND SOFTWARE AVAILABILITY

The microarray data presented here have been deposited in ArrayExpress (E-MTAB-8733).

The raw data used to prepare the figures has been deposited in Mendeley Data: https://data.mendeley.com/datasets/5gmsnv8zw9/draft?a=bc216e76-d54d-4744-98a6-5cd0cf60c8c5

