## Supplemental Figures S1-S6 and Supplemental Tables S1, S2 for "JNK signaling regulates oviposition in the malaria vector *Anopheles gambiae*"

### SUPPLEMENTARY FIGURES

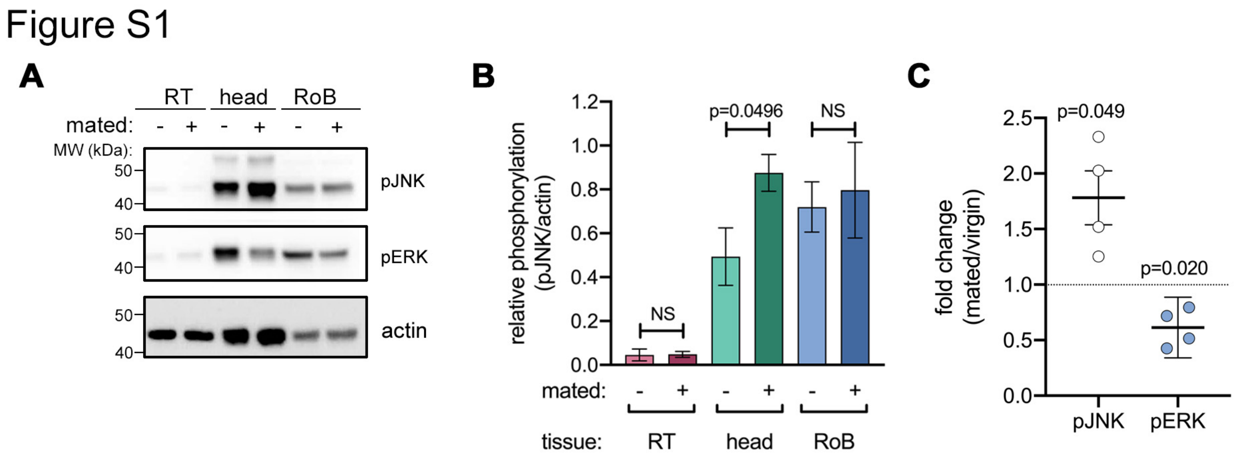


**Figure S1. Mating induces pJNK selectively in the head. A.** Representative western blot of reproductive tracts (RT; ovaries, atrium and spermatheca), heads or rest of the body (RoB) prepared from virgin (-) or mated (+) females 2 hours after mating. In **B** the optical density of bands was quantified (ImageJ) and the pJNK signal was normalized against actin and expressed as 'relative phosphorylation'. Data represent the mean ± SEM for four (RT, head) or three (RoB) independent biological replicates. Differences in the relative pJNK levels in virgin and mated tissues were analyzed using a two-tailed t test and significant *p* values ($p<0.05$) reported, ns denotes 'not significant' ($p>0.05$). In **C** the data represent the mating-induced fold change (mean ± 95% confidence interval) in actin-normalized pJNK and pERK signals in the head in four independent biological replicates. Fold-change values were compared against a hypothetical value of 1 (indicated by a dotted line) using a one sample t-test. Exact *p* values are given and $p<0.05$ was taken to be significant.

Figure S2

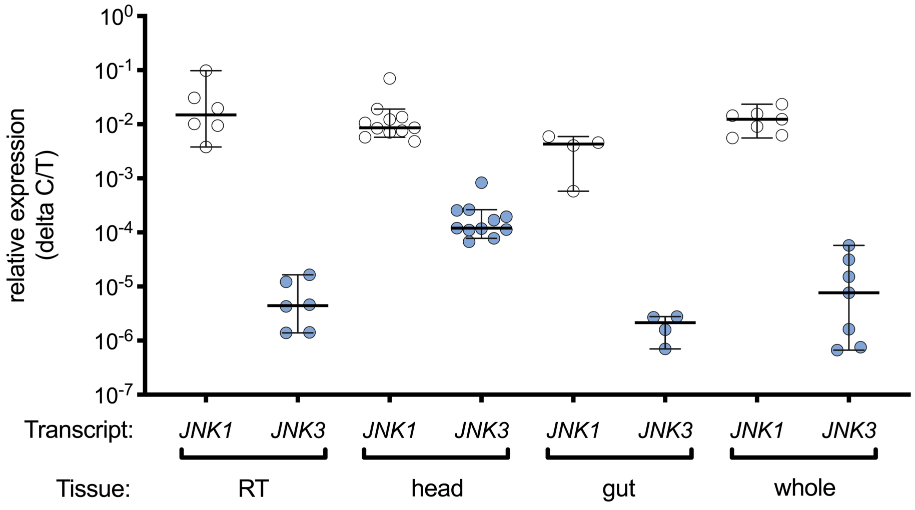


**Figure S2. Relative expression of *JNK1* and *JNK3* in *An. gambiae* tissues.** The expression of transcripts for *JNK1* (AGAP029555) and *JNK3* (AGAP009460) was measured by qRT-PCR and expressed relative to *Rpl19*, a loading control (delta C/T), in the indicated tissues or from 'whole' virgin 3-day-old females. Each point represents an independent biological replicate and bars represent the median ± 95% confidence interval of values detected.

Figure S3

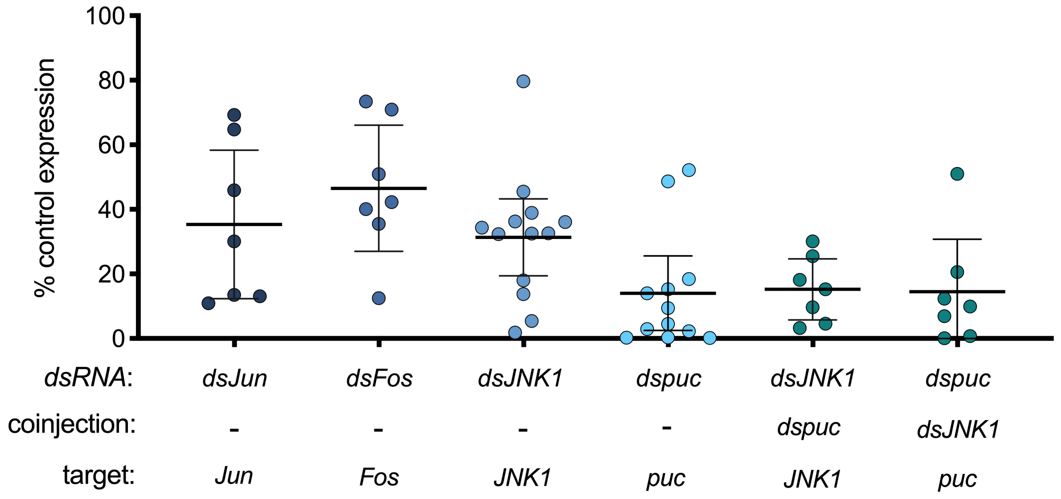


**Figure S3. Efficiency of RNAi-mediated knock down with *dsRNAs*.** The effect of single or combined *dsRNA* treatments on target gene expression was measured in whole females, two days (*JNK1, puc, joint JNK1/puc*) or five days (*Jun, Fos*) after injection. The data presented represent the % of *Rpl19*-normalized target expression in *dstarget* vs *dsGFP*-injected controls (delta delta C/T). Each dot represents an independent biological replicate and the bars represent the mean ± 95% confidence interval. Differences between *dstarget*- and *dsGFP*-treated values were analyzed using a one-sample t-test. The mean ± SEM knock down efficiencies were: *dsJun* 65±9% ($p$=0.0005, n=7); *dsFos* 53±8% ($p$=0.0005, n=7); *dsJNK1* 69±5% ($p$<0.0001, n=13), *dspuc* 86±5% ($p$<0.0001, n=12). Coinjection of *dsJNK* and *dspuc* effectively reduced expression of both *JNK1* (85±4%, $p$<0.0001, n=7) and *puc* (85±7%, $p$<0.0001, n=7).

Figure S4

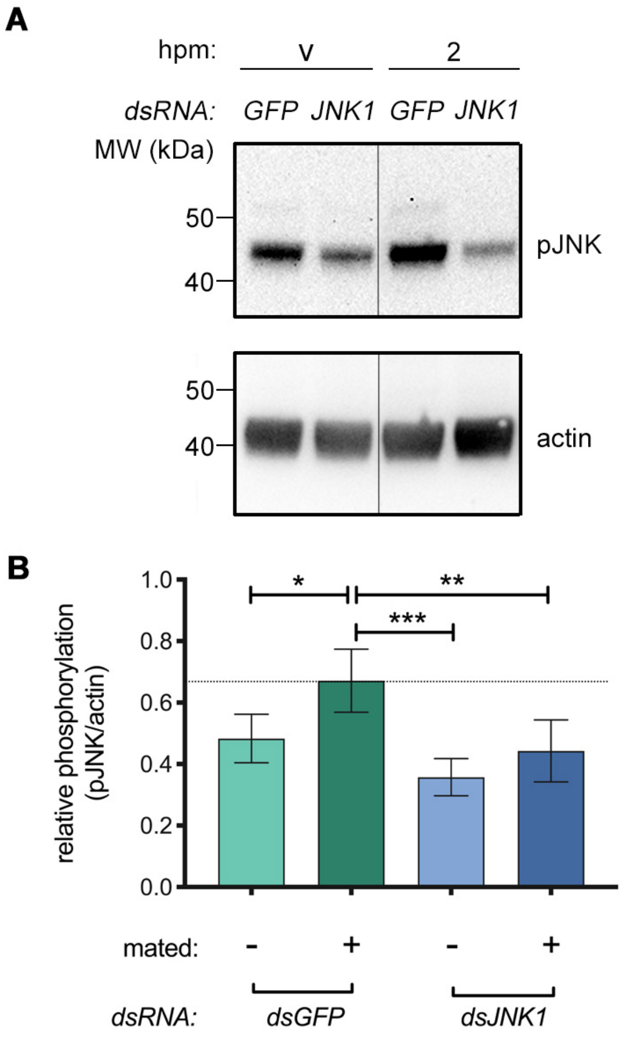


**Figure S4. *dsJNK1* inhibits mating-induced increase in pJNK in the head. A.** Representative western blot of heads dissected from *dsGFP*- or *dsJNK1*-injected virgin (v) or mated females at 2 hours post mating (hpm). Samples were western blotted using anti-pJNK then stripped and re-probed with anti-actin as loading control. The optical density of bands was quantified (ImageJ), the pJNK signal normalized against actin and expressed as 'relative phosphorylation'. The line separating the virgin and 2hpm time points indicates the removal of an irrelevant intervening lane. Data in panel **B** represent the mean ± SEM of 5 similar experiments. Differences between the mated, *dsGFP*-injected group (indicated by a dotted line) and all other groups were compared using a 2-way ANOVA test with Dunnett's multiple comparison correction. Statistically significant ($p$<0.05) differences are indicated: * denotes $p$<0.05, ** denotes $p$<0.01, *** denotes $p$<0.001.

Figure S5

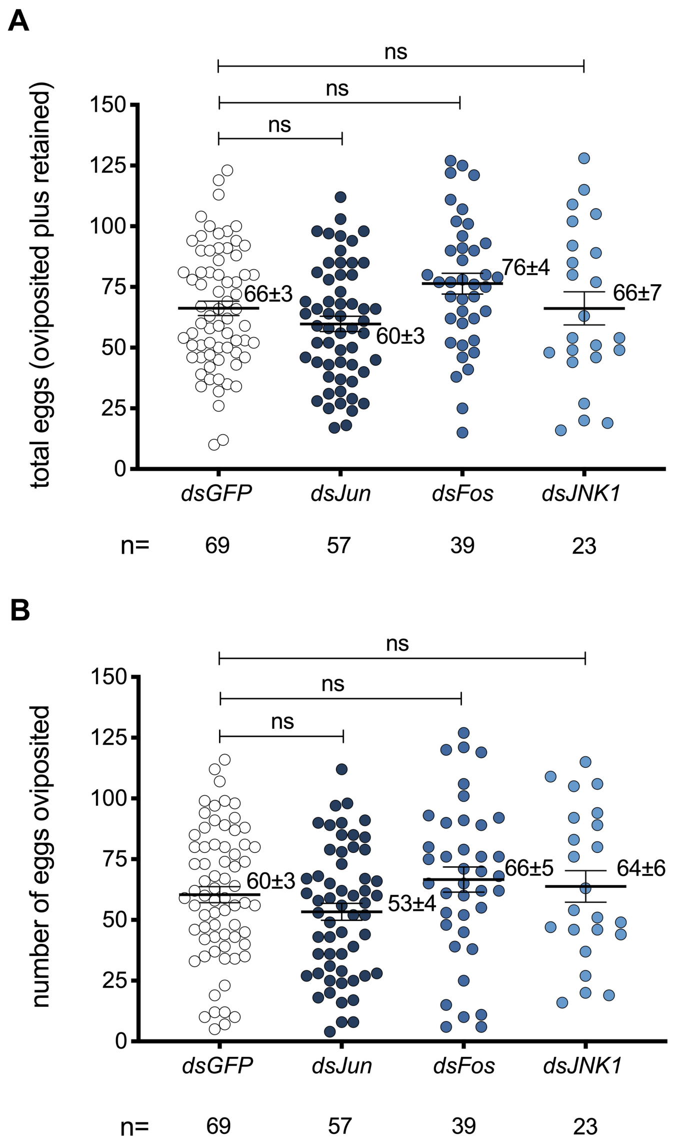


**Figure S5. JNK pathway depletion has no effect on number of eggs developed.** Virgin females were injected with the indicated *dsRNAs* and blood-fed 3 days later. After completion of egg development (2 days post blood feeding), females were mated to induce oviposition. The total number of eggs developed (the number of eggs oviposited plus the number of eggs retained in the abdominal cavity, **A**) and the number of eggs oviposited (**B**) were counted in those females who oviposited 4 days post blood feeding. The data presented represent the mean ± SEM of five, independent biological replicates. Using a one-way ANOVA test with Dunnett's multiple comparison correction all differences between control (*dsGFP*) and treatment groups were 'not significant' (ns, corrected p values >0.05).

Figure S6

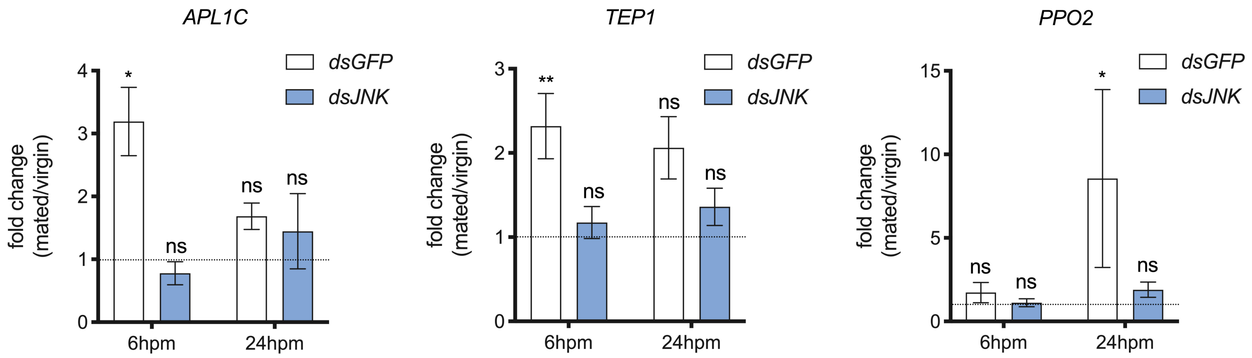


**Figure S6. Mating- and JNK1-dependence of genes expressed in the head.** Heads of *dsGFP*- (control) or *dsJNK1*- injected females were dissected at 6 and 24 hours after mating or at the same time points from age-matched virgin controls, and analyzed by qRT-PCR for the gene of interest indicated. The data presented represent the mean ± SEM fold expression change in mated females relative to age-matched virgins of the same group (1, indicated by a dotted line) and comprise at least 6 independent biological replicates. Inter-group differences at a given time point were assessed by comparing relative expression (delta CT) in the *dsGFP* virgin control group to all other treatments using a 2-way ANOVA test with Dunnett's multiple comparison correction. Statistical significance was ascribed to $p<0.05$: * denotes $p<0.05$; ** denotes $p<0.01$; ns denotes 'not significant' ($p>0.05$).

**Table S1. Low frequency of oviposition following vehicle control injection**

| Treatment | Oviposition Frequency |
|---|---|
| *dsGFP* alone (Fig. 2d) | 1.7%, n=54 |
| *dsJNK1* alone (Fig. 2d) | 3.1%, n=48 |
| *dsGFP* then 20E (Fig. 3d) | 76.2%, n=101 |
| *dsGFP* then solvent | 3.0%, n=67 |
| *dsJNK1* then solvent | 0%, n=26 |
| *dsJun* then solvent | 4.8%, n=62 |
| *dsFos* then solvent | 5.7%, n=35 |

**Table S1. Low frequency of oviposition following vehicle control injection.** Virgin females used in Figure 3c were injected with dsRNAs, blood-fed 3 days post-injection then 2 days later injected with solvent control (5% DMSO, 10% EtOH in $H_2O$) and placed in oviposition cups. The data presented represent data pooled from three or more independent biological replicates and show the number of females injected (n) and the percentage ovipositing at least one egg by day 4 post injection.

**Table S2.** Primers used in the preparation of *dsRNAs* and in qRT-PCR experiments

| **dsRNA** | **Fwd** | **Rev** |
|---|---|---|
| ***JNK1 (AGAP022950)*** | GTCACGCCTTCTACACCGTC | CAGCCCAAAGTCGAGGATT |
| ***Jun (AGAP006386)*** | CCACCCTCGAGCTGAACCTCGG | GTGGTGTTGGTGGGCACGTTCG |
| ***Fos (AGAP001093)*** | TATGATGCGCAGTGCATTTG | CGTGACTTGTGCTGATAACG |
| ***Puc (AGAP004353)*** | TGAAACATAAATCGCCCTGC | GACAGACCCTTGTAGCGCAT |
| ***GFP*** | TGTTCTGCTGGTAGTGGTCG | ACGTAAACGGCCACAAGTTC |
| ***T7*** | taatacgactcactatagggCCGCCAGTG TGCTGGAA | taatacgactcactatagggCCAGTGTGAT GGATATCTGCAGAA |
| **qRT-PCR** | | |
| ***APL1C (AGAP007033)*** | CAATGTGCTGGTTACACGCC | CCACATGTGAAGAAAATCACACTGA |
| ***PPO2 (AGAP006258)*** | GAACTTGACCCTAACGCACC | GTCCGGATACTTCTTGTCGC |
| ***TEP1 (AGAP010815)*** | CTCATGGGCTACTGGTTGT | GCTGTGAGTTAAAGTTGCTGAT |
| ***Rpl19 (AGAP004422)*** | CCAACTCGCGACAAAACATTC | ACCGGCTTCTTGATGATCAGA |
| ***Puc*** | GCCTAGTGGTCAGCTGAAGC | GCCTCGATCAGCGACAGACC |
| ***Fos*** | CAACGCTCCCGTTCAACCG | GTTGGCGTAGTACGTGTCGGC |
| ***Jun*** | AGGGCAAGTTTTGAATGCAC | CACGCACTTTCTCCCTTTGT |
| ***JNK1*** | GCCGAAGAACGACAAACTATGTGC | GCTGTGTTACTGTATCGTATGCG |

**Table S2. *dsRNA* and qRT-PCR primers used in this study.** All qRT-PCR primers were used at 300nM except *Rpl19* Rev which was used at 900nM. PCR products for dsRNA generation were all prepared using the listed primers at 200nM.
